# Functional CENP-B boxes are selectively conserved at centromeric regions

**DOI:** 10.64898/2026.05.25.727640

**Authors:** Chuan Jiao, Nikolai Goncharov, Anna Shmakova, Camellia Chakraborty, Daniele Fachinetti

## Abstract

Human centromeres are built over large stretches of repetitive, divergent α-satellite DNA. Within these sequences lies a conserved, defined 17-bp sequence named the CENP-B box that is bound by the DNA-binding protein CENP-B. Recent studies have proposed that CENP-B box motifs along chromosome arms exist and may represent conserved, ectopic binding sites with functional relevance. Here, we evaluate the genomic distribution, conservation, and binding capacity of different CENP-B box motifs outside canonical centromeres. Analysis of thousands of complete human centromere assemblies reveals exceptional conservation of a unique and canonical CENP-B box motif within centromeres. This is in contrast with the high sequence variability and stochastic occurrence of more degenerate CENP-B box motifs along chromosome arms. Consistently, CENP-B binds only at canonical CENP-B box motifs embedded within α-satellite sites, with no evidence of functional binding at any ectopic sites. Together, these results indicate an adaptive selection of canonical CENP-B boxes within centromeric regions, in contrast to random sequence occurrences for ectopic CENP-B box-like motifs.

## Introduction

Centromeres are essential chromosomal elements that ensure accurate chromosome segregation across eukaryotes. In humans, centromere position is primarily specified by epigenetic mechanisms (McKinley and Cheeseman 2016; Mellone and Fachinetti 2021), yet canonical human centromeres are assembled over large stretches of tandemly repeated DNA arrays, named α-satellite DNA (Manuelidis and Wu 1978; Waye and Willard 1985; Talbert and Henikoff 2022). Centromeric α-satellite is built from AT-rich ∼171-bp monomers that can be organized into chromosome-specific <u>H</u>igher-<u>O</u>rder <u>R</u>epeats (HORs). Multimeric repeat units that are themselves tandemly repeated to form megabase-scale arrays, often transitioning at their edges into more diverged/less-ordered monomeric α-satellite and other pericentromeric repeats (Willard and Waye 1987; Miga et al. 2020; Logsdon et al. 2021; Altemose et al. 2022). Recent telomere-to-telomere assemblies and centromere maps show that these arrays are highly variable in length, sequence composition, and internal layering across chromosomes and between haplotypes/individuals, while still providing a common substrate on which the kinetochore assembles (Altemose et al. 2022; Logsdon et al. 2025; Hansen et al. 2025; Volpe et al. 2025; Gao et al. 2025).

Within these repetitive α-satellite arrays resides a short and conserved 17-bp DNA motif known as the CENP-B box, which serves as the binding site for the DNA-binding centromere protein B (CENP-B) (Masumoto et al. 1989). In human centromeres, CENP-B boxes are typically positioned within every second α-satellite monomer, resulting in an approximate ∼340-bp periodicity corresponding to the HOR organization (Henikoff et al. 2015). However, this regular spacing is not universal, as certain centromeric arrays display disrupted or heterogeneous CENP-B box organization (Corda and Giunta 2025) or even a complete absence of the motif, as in the α-satellite HOR array on the human Y chromosome (Miga et al. 2014; Henikoff et al. 2015). CENP-B binds to the CENP-B box through its N-terminal DNA-binding domain, which recognizes a “canonical motif” 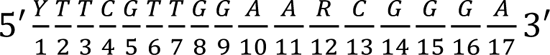 with degenerate bases at positions 1 (Y = C/T) and 12 (R = A/G)(Masumoto et al. 1989; Yoda et al. 1992; Muro et al. 1992). Mutational *in vitro* analyses and structural studies demonstrated that CENP-B specificity to the 17-bp motif is not uniformly distributed (Sugimoto et al. 1997, 1998; Ohzeki et al. 2002; Tanaka et al. 2001). Here it was shown that protein binding depends on two short sequence elements at positions 2-5 (TTCG) and 13-15 (CGG), which together form the primary DNA-contact interface with CENP-B. Nucleotide substitutions at positions 1 and 17 did not significantly influence binding, whereas substitutions at positions 3, 4, and 16 did. This infers that the internal 14-bp motif 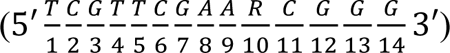 constitutes the minimum requirement for CENP-B binding. Nevertheless, this shorter motif exhibited lower affinity for CENP-B compared to the 17-bp motif. The inclusion of a T at the 5′ end and the A or T at the 3′ end 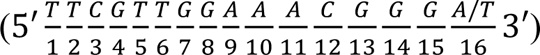 resulted in an affinity comparable to that of the authentic motif. The relevance of the A at the 3′ end was further demonstrated by structural studies, along with a synthetic higher-order α-satellite experiment that also demonstrated the importance of the central A in position 10 (Okada et al. 2007; Tanaka et al. 2001).

Partially consistent with biochemical evidence demonstrating tolerance to substitution at specific positions within the CENP-B box, the “broad motif” is commonly represented as 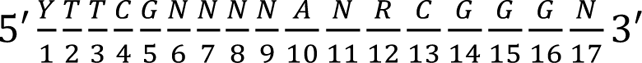 However, genome-wide analyses often employ an even more “degenerated motif” 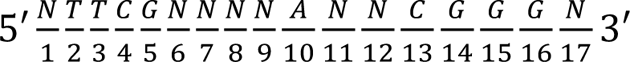 to account for sequence divergence accumulated through the rapid evolution of α-satellite arrays (Altemose et al. 2022). Using this “degenerated motif”, *Corda and Giunta* identified several dispersed CENP-B boxes outside canonical centromeric arrays (Corda and Giunta 2025). In their study, the authors proposed functional implications of these ectopic motifs, possibly influencing chromatin structure via long-range interactions due to CENP-B binding and likely dimerization.

In our study, we aim to assess whether the ectopic CENP-B boxes represent functionally meaningful binding sites for CENP-B or if instead they reflect neutral sequence variation present outside centromeric chromatin. To address this question, we directly compared the genomic distribution, positional enrichment, and sequence conservation of canonical, broad, and degenerated motifs in the CHM13 (Nurk et al. 2022), RPE-1 (Volpe et al. 2025) and HG002 (Liao et al. 2023; Hansen et al. 2025) full genome assemblies, as well as across 2,110 complete human centromere assemblies (Gao et al. 2025) and across related chromosome arm contigs from the Human Pangenome Reference Consortium (HPRC) dataset. To address the functional importance, we analyzed CENP-B binding across the genome along with transcriptomics data of cells overexpressing CENP-B.

## Results

### The canonical CENP-B box motif is only conserved within centromeres, while ectopic CENP-B box motifs reflect neutral sequence variation

To identify all potential CENP-B binding motifs across both centromeric regions and chromosome arms, we first analyzed three complete human genome assemblies: CHM13, RPE-1, and HG002 (Nurk et al. 2022; Volpe et al. 2025; Liao et al. 2023; Hansen et al. 2025). Restricting motif searches risks underestimating motif abundance in the genome. Conversely, relaxing the motif, allowing mismatches, increases sensitivity but raises the possibility of detecting sequences with no functional relevance. Accordingly, we mapped the orientation and genomic position of all CENP-B boxes across the three reference genomes (**Figure 1A,B, S1A,B** and **S2A,B**) based on their consensus definition (canonical, broad or degenerated motif as in **Table S1**). As expected, we detected a strong enrichment of all three types of CENP-B box motifs within centromeric regions across all examined genomes, underscoring the selective maintenance of CENP-B boxes at human centromeres (**Figure 1A-C, S1A-C, S2A-C** and **Table S2**). Consistent with *Corda and Giunta* (Corda and Giunta 2025), we observed a minor occurrence of ectopic CENP-B box motifs along chromosome arms for the degenerated consensus motif (∼0.44–0.5% of all CENP-B boxes), whereas the broader motif showed a markedly lower arm density (∼0.2%) (**Figure 1C, S1C, S2C** and **Table S2**). It was extremely rare to find the canonical motif outside centromeres (∼0.01%), indicating strong sequence constraint.

**Figure 1.**
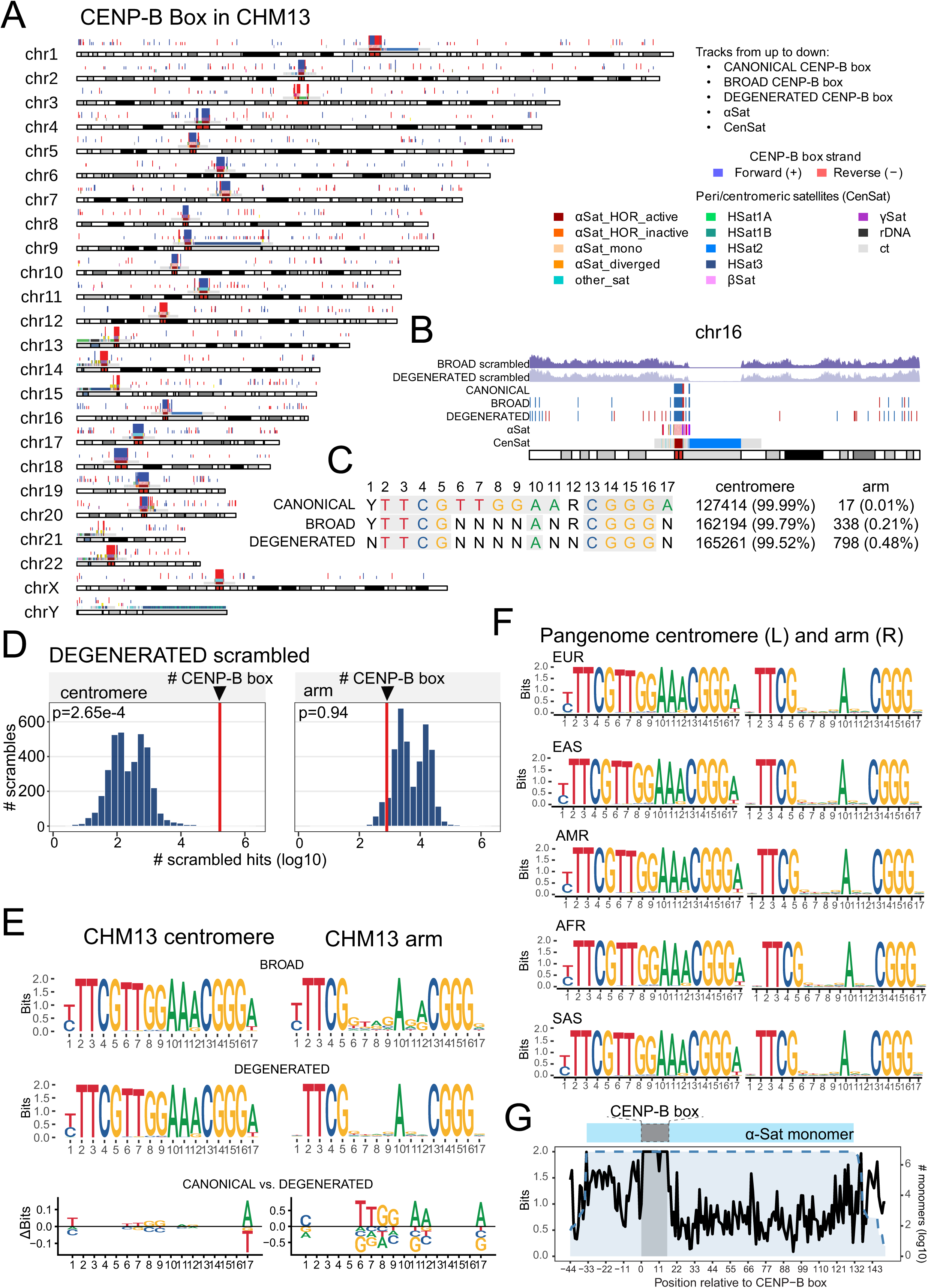
CENP-B box motif is selectively conserved at centromeres, whereas arm-located CENP-B box motifs are stochastic. **(A)** Karyoplot of CENP-B box motifs in the CHM13 reference genome. The first three tracks show CENP-B box motifs based on canonical, broad, and degenerated definitions, with forward- and reverse-strand occurrences shown in blue and red, respectively. The fourth and fifth tracks show α-satellite (αSat) monomer classes and (peri-)centromeric satellites classes, respectively, colored by annotation class. (Peri-)centromeric satellite classes include active and inactive αSat higher-order repeats (HORs), diverged αSat, and other satellite families such as HSat1–HSat3, β-satellite (βSat), γ-satellite (γSat), rDNA, and centromeric transition (ct) regions; see color key. **(B)** Representative chromosome 16 from panel A. The two upper tracks show the density (number of hits per 100kb window) of scrambled motif matches for the degenerated and broad CENP-B box, respectively. Below these tracks, observed CENP-B box motifs are shown as strand-specific ticks, followed by annotation tracks for αSat and centromeric satellites classes. **(C)** Definitions of canonical, broad and degenerate CENP-B box motifs and distributions in the CHM13 reference genome. **(D)** Comparison between observed CENP-B box motif counts and scrambled motif counts in CHM13 centromeric regions and chromosome arms. Histograms show the null distributions generated from scrambled motifs, with the observed CENP-B box count indicated by a red vertical line. p-values represent empirical one-sided permutation probabilities, defined as the proportion of scrambled sequences yielding motif counts greater than the observed counts. **(E)** Sequence logos of broad and degenerated CENP-B box motifs in CHM13 centromeric regions (left) and chromosome arms (right). Bits (0-2) represent the information content at each motif position, with higher values indicating greater nucleotide conservation. ΔBits plots highlight positional differences between canonical and degenerated motifs. **(F)** Sequence logos of CENP-B box motifs generated from centromere (left) and arms (right) across continental population groups (African (AFR), European (EUR), East Asian (EAS), South Asian (SAS), and American (AMR). **(G)** Sequence conservation of αSat monomers across continental population groups. The CENP-B box is highlighted in grey.

To directly test whether ectopic motifs reflect random sequence composition, we compared the observed motif frequencies with all possible combinations of 17 bp scrambled control sequences generated by randomly shuffling the fixed nucleotides of the degenerated CENP-B box motifs (**Figure 1B** and **Table S3**). We compared centromere and arm motif counts with a null distribution generated by scrambling the same sequences. If motif occurrence is driven by sequence composition, counts in real and scrambled sequences should scale together, while deviations from this distribution indicate sequence-specific selective enrichment. Consistent with this framework, degenerated motifs within centromeric regions showed a clear deviation from random expectation: the observed number of matches was significantly higher than predicted by the null model, falling in the extreme tail of the scrambled distribution (p = 2.65 × 10^-4^) (**Figure 1D**). In contrast, motif occurrence in chromosome arms followed the expectation for random sequence composition, in which the observed counts of degenerated motifs were not significantly different from the scrambled distribution (p = 0.20; **Figure 1D**). Together, these results indicate that while degenerated CENP-B box motifs are selectively enriched in centromeric regions, their presence in chromosome arms is largely explained by random sequence composition, consistent with the behavior expected for short, degenerate motifs. Same results were obtained in RPE-1 and HG002 (**Figure S1D** and **S2D**).

To investigate the selective pressure of maintaining a unique nucleotide motif for CENP-B binding, we performed a sequence variation analysis in CHM13, RPE-1 and HG002. Here, we revealed striking differences between centromeric and ectopic motifs. Within centromere-spanning α-satellite arrays across all chromosomes, CENP-B boxes displayed strong nucleotide positional conservation matching the canonical motif, regardless of whether broad or degenerated motif definitions were used for analysis (**Figure 1E** and **S3**). In contrast, motifs detected along chromosome arms exhibited substantially higher nucleotide variability at all nucleotides not fixed in the degenerated motif (**Figure 1E** and **S3**). Similar results were observed in the analysis of 2,110 haplotypes assemblies from 65 individuals (Gao et al. 2025), representing diverse continental ancestry groups (**Figure 1F**).

Across this dataset, the canonical CENP-B box sequence showed high nucleotide conservation, with no detectable variation across African (AFR), European (EUR), East Asian (EAS), South Asian (SAS), and American (AMR) populations. Importantly, this pattern of conservation was preserved when progressively relaxed motif definitions were applied, indicating that the observed invariance is not an artefact of motif stringency. On the contrary, ectopic CENP-B box motifs detected within the chromosome arms exhibited higher nucleotide variability (**Figure 1F**). Extending the sequence variation analysis to the whole α-satellite monomers containing CENP-B boxes within the 2,110 haplotypes, we reveal that the CENP-B box itself remains highly conserved within human centromeres, despite the extensive structural and sequence variation that is characteristic of α-satellite higher-order repeat arrays (**Figure 1G**). These results point to strong and specific selective pressure acting on the CENP-B box motif within centromeres, maintaining sequence integrity despite rapid evolution and diversification of the surrounding α-satellite DNA.

### No evidence of functional CENP-B binding in chromosome arms

We then investigated in detail the location of the CENP-B box motifs that were found in the chromosome arms. Unlike the broad or degenerated motifs, all the canonical motifs that identified along chromosome arms co-localized with clusters of ectopic degenerate α-satellite sequences (**Figure 2A**). These regions were previously reported on the arms of chromosomes 2 and 4 in the CHM13 reference (**Figure 2B** and **S4**) (Altemose et al. 2022). Therefore, these canonical ectopic CENP-B box motifs likely originated from displaced or translocated centromeric satellite DNA rather than being formed *de novo* as new binding sites for CENP-B. On the contrary, the ectopic broad and degenerated CENP-B box motifs are randomly distributed along the chromosome arms without any association with α-satellite DNA.

**Figure 2.**
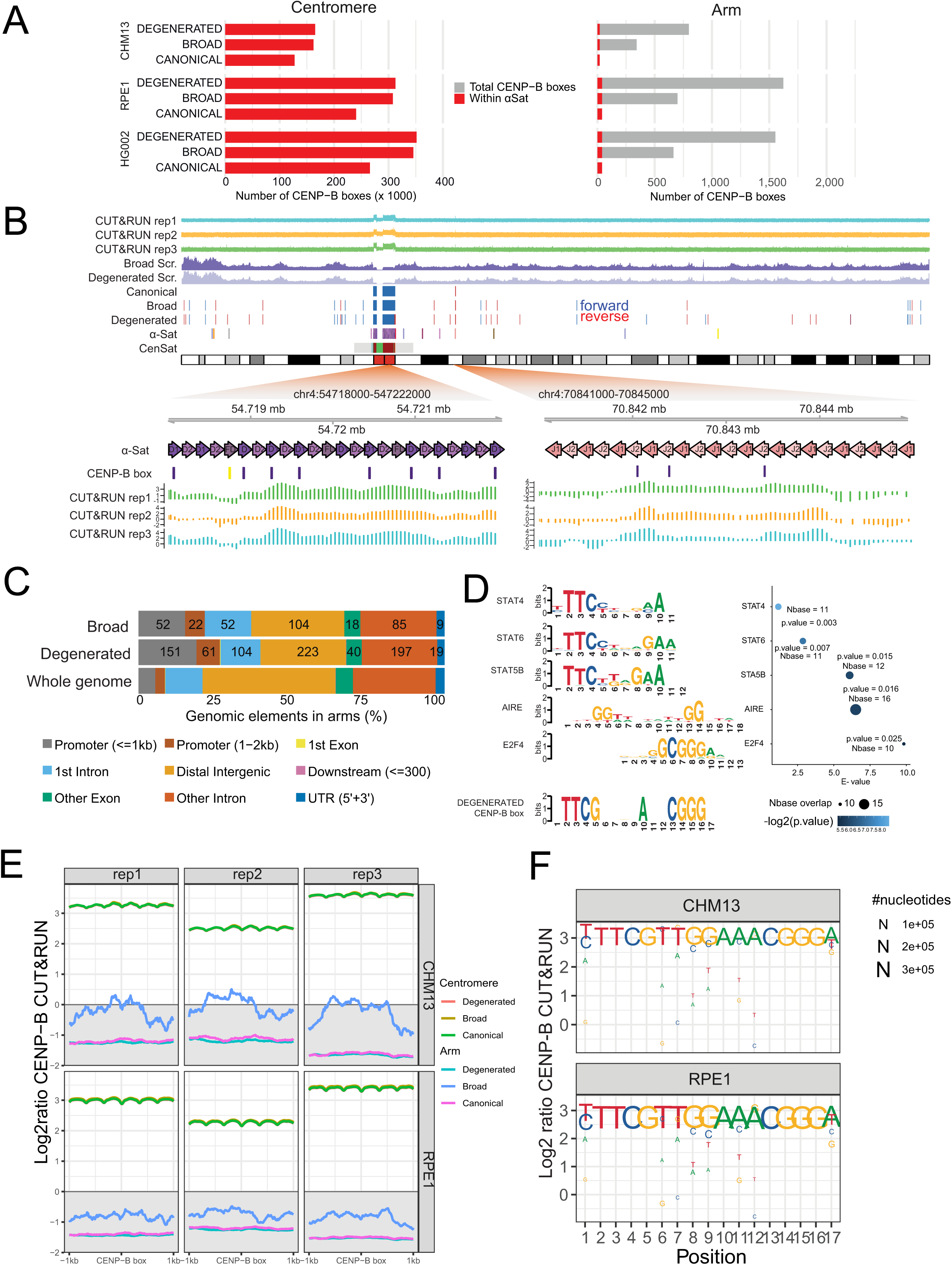
CENP-B occupancy is preferentially associated with motifs that retain the canonical sequence motif. **(A)** Genome-wide distribution of CENP-B box motifs across the human genome assemblies CHM13, RPE-1, and HG002. Bar plots show the number of CENP-B boxes and the proportion located within αSat arrays (red) relative to the total number of motifs (grey), stratified by canonical, broad and degenerated motif classes and separated into centromeric and chromosome arm regions. **(B)** Centromere and chromosome arm CENP-B box clusters located on chromosome 4 displaying CENP-B CUT&RUN signal when mapped to the CHM13 genome reference. Tracks show genomic coordinates, motif orientation (forward in blue, reverse in red), scrambled motif density, and CUT&RUN signal across three biological replicates. These loci coincide with blocks of αSat-like sequence. Yellow (degenerated) and violet (canonical) marking shows distinct CENP-B box motifs. **(C)** Genomic annotation of broad and degenerated CENP-B box motifs on chromosome arms. Bars show the proportion of motif occurrences overlapping annotated gene elements in chromosome arms, compared with their proportion in the whole genome. The actual number of CENP-B box motifs in the respective categories is indicated. **(D)** Motif similarity between the degenerated CENP-B box and known human transcription factor binding motifs. Sequence logos of the top-scoring matches (STAT4, STAT6, STAT5B, AIRE, and E2F4) are shown alongside the degenerated CENP-B box query motif. The dot plot summarizes Tomtom similarity statistics, including p-values (probability of observing a match of equal or greater similarity for a single comparison) and alignment overlap. E-values indicate the expected number of such matches occurring by chance across the database search. **(E)** Aggregate CUT&RUN signal profiles centered on CENP-B box motifs with 1 kb flanking intervals in CHM13 and RPE-1 for canonical, broad, and degenerated motifs in centromeric and chromosome arm regions. **(F)** Nucleotide-specific CENP-B CUT&RUN enrichment across the 17-bp motif in CHM13 and RPE-1 reference genomes. Position-specific log2 enrichment values (CUT&RUN signal relative to IgG control) are shown for each nucleotide (A, C, G, T) at each motif position, calculated from motif instances overlapping CUT&RUN signal.

Consistent with previous reports (Corda and Giunta 2025), a small subset of chromosome arm CENP-B-like motifs in the broad (21.6%) and degenerated (26.6%) configurations were enriched (∼3 fold more compared to whole genome) within a 2 kb window around annotated transcription start sites (TSS) (**Figure 2C**). To analyse the regulatory context of the CENP-B box motifs detected on chromosome arms, we intersected these loci with ENCODE candidate cis-regulatory elements (cCREs) (**Figure S5**). This analysis revealed an enrichment in promoter- and enhancer-associated annotations, with the strongest enrichment observed for promoter-like signatures (Benjamini–Hochberg adjusted p-value <0.05), including promoter-like elements (fold enrichment = 22.98), proximal enhancer-like signatures (10.23), and accessible H3K4me3 (6.93), followed by distal enhancer-like elements (2.73) and transcription factor-associated regions (2.16). In contrast, chromatin-accessible elements without promoter-associated signatures showed weak or no enrichment, and accessible CTCF-associated or chromatin-accessible regions with transcription factor binding evidence (CA–TF classes) were not enriched.

To determine whether the degenerated CENP-B box motif resembles known transcription factor binding sites, we compared the degenerated motif against the human transcription factor motif database using Tomtom (MEME Suite), which evaluates motif similarity (Gupta et al. 2007). This comparison identified five transcription factor motifs with partial similarity to the degenerated CENP-B box motif, including STAT4 (p = 0.003, E = 0.11), STAT6 (p = 0.001, E = 0.03), STAT5B (p = 0.001, E = 0.015), and, with lower similarity, AIRE (p = 0.016, E = 0.36) and E2F4 (p = 0.025, E = 0.10) (**Figure 2D**). Here the p-value reflects the probability of observing a motif match with equal or greater similarity by chance for one motif comparison, while the E-value represents the expected number of matches with equal or greater similarity that would occur by chance across the entire database search. Thus, lower E-values reflect stronger similarity, while values approaching or exceeding 1 indicate matches that are likely to occur frequently by chance. The relatively modest E-values observed here indicate that similar matches are expected to occur by chance when comparing the query motif to any transcription factor motifs in the database. Consistent with this interpretation, the overlap between the degenerated CENP-B box and the identified transcription factor motifs is restricted to short sequence fragments, rather than reconstructing the full 17-bp canonical CENP-B box motif. Thus, these motif alignments reflect local sequence resemblance rather than shared motif between CENP-B box and transcription factor binding sites. We then analyzed if CENP-B could influence the regulation of the 202 genes containing CENP-B box motifs within the +/– 2 kb of their TSS. First, we observed that the distribution of the motif was mostly heterogeneous along the 4 kb window (**Figure S6**), suggesting no consistent functional relationship between the motif’s position and gene regulation. Second, we analyzed the transcriptome of human foreskin fibroblasts overexpressing a DOX-inducible CENP-B construct (**Figure S7A**). Gene set enrichment analysis of a custom gene set comprising genes with CENP-B-box motif in the +/– 2 kb window of the TSS (116 genes detected in RNA-sequencing experiment) revealed no significant enrichment upon CENP-B overexpression, indicating that CENP-B does not exert a consistent transcriptional activation or repression effect on these genes (**Figure S7B**). Altogether, these results imply that, despite having potential binding sites, CENP-B is unlikely to directly regulate transcription of these genes either because it does not bind to these ectopic sites or because it is unable to affect gene expression.

To investigate further these possibilities, we assessed the binding capacity of CENP-B at the degenerated motifs detected along the chromosome arms. We mapped on CHM13 and RPE-1 references genome-wide CENP-B CUT&RUN experiments previously performed on RPE-1 cells (Dumont et al. 2020) following normalization to the IgG control. As expected, aggregate profiles centered on CENP-B box motif positions revealed robust enrichment at centromeric sites across all three biological replicates (log2 enrichment values of > 2 over background; **Figure 2E**). Accordingly, CENP-B CUT&RUN signal was strongly enriched at the centromeric regions, representing >99.9% of all overlapping peaks when mapped to CHM13 (**Figure S8**). In contrast, CENP-B-like motifs located along chromosome arms showed no reproducible CENP-B enrichment, with signal levels remaining close to background (log2 ratio ≈0) irrespective of motif definitions (**Figure 2E** and **S8**) except for three ectopic CENP-B peaks detected in all 3 replicates. These three peaks, however, were detected in chromosome 4 and 2 ectopic sites where CENP-B boxes were embedded within degenerate α-satellite arrays (**Figure 2B** and **S9**). It is important to note that, given the high sequence similarity of centromeric α-satellite regions and the short-read nature of CUT&RUN, reads are prone to multi-mapping and preferential assignment to the best-scoring reference locus, so reported CENP-B occupancy outside centromeres should be interpreted cautiously. Mapping the same CUT&RUN dataset to the RPE-1 reference, the same cell line from which the CUT&RUN data was generated (Dumont et al. 2020), eliminated these arm-associated signals (**Figure S8**). This suggests that their detection in CHM13 reflects a reference-dependent mapping artifact rather than *bona fide* ectopic CENP-B occupancy.

It was recently reported the presence of few hundreds (379) of CENP-B peaks on chromosome arms enriched at regions of negatively supercoiled DNA containing repetitive sequences, such as multiple CCAAT boxes, but not at ectopic CENP-B-boxes (Wu et al. 2026). Here the authors used a combination of low/high salt CUT&RUN method in RPE-1 cells mapped on CHM13. We therefore extend our peaks analysis to the chromosome arms outside the CENP-B-box motif using our high salt CUT&RUN analysis (Dumont et al. 2020). Regardless of mapping to CHM13 or RPE-1 genome, we detected only few CENP-B peaks in chromosome arms mostly (if not all) located at the periphery of centromeric regions (**Figure S10A**), despite higher coverage and centromeric enrichment of our dataset compared to theirs (Wu et al. 2026) (**Figure S10B,C**).

These results suggest that while transient binging of CENP-B to other motifs could occasionally occur –expected for a DNA binding protein –stable binding occurs only to CENP-B boxes within centromeric regions. Interestingly, quantitative comparison of the distinct classes of CENP-B box motifs and CENP-B CUT&RUN peaks revealed a clear relationship between motif sequence integrity and protein binding. By summarizing CENP-B box sequence variants present at CENP-B binding sites, we observed that CUT&RUN peaks predominantly colocalized with motifs closely matching the canonical CENP-B box sequence rather than with broad or degenerated variants (**Figure 2F** and **Table S4**). CUT&RUN enrichment values were also compared across genomic motif instances grouped by motif class (**Figure S11)**. At centromeres, enrichment differed significantly among canonical, broad and degenerated motifs, with canonical motifs showing the highest signal. Pairwise comparisons showed significantly higher enrichment at centromeric canonical motifs than at centromeric broad and degenerated motifs (adjusted p = 3.1 × 10⁻⁵ and 4.4 × 10⁻¹⁴, respectively). In chromosome arms, motif-associated CUT&RUN signal remained weak overall, with median values close to background. Detectable arm-associated signal was limited to a small number of α-satellite–embedded loci on chromosomes 2 and 4 that retain canonical CENP-B boxes. Together, these results indicate that sequence integrity of the CENP-B box is a primary determinant of CENP-B binding specificity, with canonical motifs representing the major high-affinity binding sites across the genome.

## Discussion

Historically, genome-wide analyses of putative CENP-B binding sites have adopted degenerate representations of the canonical CENP-B box such as the broad motif (Sugimoto et al. 1998) and, later, the more permissive degenerated motif (Altemose et al. 2022). This strategy was conceptually justified by the biochemical evidence demonstrating tolerance to nucleotide substitution at specific positions within the 17-bp motif (Sugimoto et al. 1997, 1998; Tanaka et al. 2001; Ohzeki et al. 2002) and by the rapid evolution of α-satellite DNA, which exhibits substantial sequence divergence. However, this reasoning assumes that the observed variability in the α-satellite arrays extends uniformly to the CENP-B box itself. Our sequence variation analyses challenge this assumption as we find that the canonical CENP-B box motif is extraordinarily conserved across centromeric arrays, not only across reference assemblies (CHM13v2.0, RPE-1v1.1, HG002v1.1) but also across 2,110 fully assembled centromeres from 65 individuals representing deeply divergent human haplotypes (Gao et al. 2025). This conservation stands in sharp contrast to the surrounding α-satellite sequence, which shows extensive polymorphisms in monomer composition, HOR structure, and array length. Our observations indicate that, while biochemical assays demonstrate positional tolerance under controlled *in vitro* conditions, the *in vivo* evolutionary realization of functional CENP-B boxes is subject to exceptionally strong selective constraints. Accordingly, CENP-B binds to canonical CENP-B box motifs within the centromeric regions rather than on more degenerated motifs, suggesting selective pressure to maintain those unique motifs, potentially arising from interactions with additional protein factors or from structural constraints imposed by centromeric chromatin organization. Sequence homology links CENP-B to pogo-like DNA transposases, supporting its origin via transposase domestication within the host genome, with loss of endonuclease activity and retention of sequence-specific DNA binding to the CENP-B box (Mateo and González 2014; Casola et al. 2007). Based on our results, we speculate that the transposase-derived CENP-B–like proteins were first recruited to centromeric regions, likely in the context of transposon control, and only later the specific high-affinity binding sites (CENP-B boxes) became fixed and/or amplified in some satellites that benefited from tighter CENP-B recruitment (Gamba and Fachinetti 2020). However, it remains unclear why in certain equids, CENP-B boxes are found in older satellites that have lost centromeric function, whereas the CENP-A–bound core repeats lack those motifs and CENP-B (Cappelletti et al. 2025).

The finding that CENP-B box motifs exist outside centromeres in the human genome (Corda and Giunta 2025) is compelling considering the multiple other known roles of CENP-B besides being a key centromeric protein (Suzuki et al. 2004; Hoffmann et al. 2016; Fachinetti et al. 2013, 2015; Dumont et al. 2020) such as acting as an epigenetic regulator (Okada et al. 2007; Morozov et al. 2017; Otake et al. 2020), favoring nucleosome repositioning (Nagpal et al. 2023), regulating genome architecture (Sugimoto et al. 1994; Yoda et al. 1998; Tanaka et al. 2001; Kasinathan and Henikoff 2018; Chardon et al. 2022) and transcription (Chen et al. 2021; Ishikura et al. 2021). However, our data suggests that on chromosome arms degenerative CENP-B boxes have stochastic sequence composition rather that adaptive selection. This explains why numerous CENP-B box-like motifs can be identified outside canonical centromeric arrays. We also show that these chromosome arm sites lack evidence of functional binding, as no consistent CENP-B peaks were detected. Our data do not exclude that CENP-B could transiently bind at these ectopic sites or other regions, like negatively supercoiled DNA containing repetitive sequences, such as CCAAT boxes (Wu et al. 2026) in a manner that is undetectable by high salt CUT&RUN, but it remains unclear what functional role such a short-lived interaction might play. We have previously shown that CENP-B could form DNA loops via its dimerization domain by binding two CENP-B boxes that are in proximity to each other (∼ 1 kb or less(Chardon et al. 2022)). The presence of only ∼1000 total sites in the degenerated conformation distributed across genome with an average distance of 8.2 Mb and with only 17 of these sites at a distance in the range of 1 kb (**Table S5**) argues against any potential functional role for CENP-B dimerization in chromosome organization outside centromeres.

CENP-B was proposed to directly or indirectly regulate transcription (Chen et al. 2021; Ishikura et al. 2021). However, this was demonstrated in the context of repetitive DNA where many CENP-B boxes are present, in contrast to a sporadic binding at a few consecutive sites around TSS. The higher incidence of CENP-B box-like motifs could be indeed stochastic or, to a certain degree, due to sequence similarity to transcription factor binding sites. Functional experiments using gene knock-out or by deleting these binding sites might be required to formally assess any potential regulatory consequences. In this regard, we did not detect any consistent changes in the transcription level of genes harboring CENP-B box motifs within their TSS upon CENP-B overexpression, suggesting that transient CENP-B binding to those sites is unlikely to have any functional direct effect. In the same line, the findings that gene regulation of certain genes is affected in the absence of CENP-B, regardless of whether CENP-B is found at these promoters (Wu et al. 2026), strongly suggest for an indirect effect of CENP-B alteration on gene transcription. For example, the mild upregulation of replication-coupled histone / nucleosome genes observed in the context of CENP-B depletion might be due to changes in the heterochromatin content(Okada et al. 2007) or to defects in centromere replication (Salinas-Luypaert et al. 2025; Scelfo et al. 2024). Finally, the epigenetics status of these putative binding sites add another level of regulation for CENP-B binding as in the case of DNA methylation (Tanaka et al. 2005; Salinas-Luypaert et al. 2025).

Taken together, our findings demonstrate that only the canonical CENP-B box embedded within α-satellite DNA displays evolutionary conservation in humans and binding competence characteristic of functional CENP-B recognition sites. In contrast, CENP-B box-like motifs located along chromosome arms are highly heterogeneous, sporadically distributed, and lack detectable CENP-B binding activity, like other regions on chromosome arms. These results support a model in which ectopic CENP-B box-like sequences arise through random sequence variation rather than selective pressure, underscoring the unique specificity of CENP-B for its authentic centromeric binding motif. The usage of degenerate CENP-B box motifs, while methodologically appropriate for exploratory surveys of satellite-derived sequences, should not be interpreted as evidence of functional CENP-B boxes (i.e. bound by CENP-B). Our findings highlight that biochemical tolerance to nucleotide substitution *in vitro* does not necessarily translate to a widespread sequence variability *in vivo*.

## Materials and Methods

### Identification of centromeric regions and chromosome arms

Genome reference FASTA files and RepeatMasker annotations were obtained for CHM13v2.0 (Nurk et al. 2022), RPE-1v1.1 (Volpe et al. 2025) and HG002v1.1 (Liao et al. 2023; Hansen et al. 2025). To define (peri-)centromeric intervals consistently across the three assemblies, we used the centromereAnnotation.wdl workflow from alphaAnnotation (https://github.com/kmiga/alphaAnnotation) together with annotation files rDNA1.0.hmm, AS-HORs-hmmer3.4-071024.hmm, and AS-SFs-hmmer3.0.290621.hmm. (Peri-)centromere intervals, alphaSat classifications and orientations, HOR intervals were identified. (Peri-)centromeric boundaries in RPE-1v1.1 and HG002v1.1 were cross-checked with CHM13v2.0 for consistency based on the repeat elements annotated at the corresponding boundary positions.

### Detection of CENP-B box in reference genomes

Whole-genome motif searches were performed in CHM13v2.0, RPE-1v1.1, and HG002v1.1 using three CENP-B box consensus definitions:

Canonical 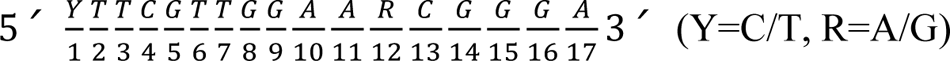

Broad 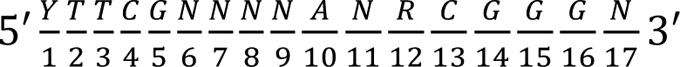

Degenerated 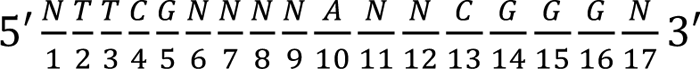

Because RPE-1v1.1 and HG002v1.1 reference genomes contain unresolved positions, motif matches were retained only if they contained at most 6 Ns for the broad motif, at most 8 Ns for the degenerated motif, and no Ns for the canonical motif. Motifs were then classified into centromere and arm region based on their (peri-)centromeric intervals.

### Detection of CENP-B box in Human Pangenome

In addition to three reference genomes, CENP-B box motifs were also detected in the human Pangenome to evaluate sequence conservation across diverse human haplotypes.

For centromeric analyses, 2110 (peri-)centromere sequences and annotations were downloaded from figshare (https://figshare.com/s/cb839beea6ed06d641db)(Gao et al. 2025).

For chromosome-arm analyses, haplotype-resolved human genome assemblies were obtained from the Human Pangenome Reference Consortium (HPRC) Year 1 release (https://github.com/human-pangenomics/HPP_Year1_Assemblies) and aligned to the CHM13v2.0 reference genome using minimap2 (v2.30)(Li 2018). To restrict analysis to chromosome arms, alignments overlapping (peri-)centromeric regions were removed.

Motif matches were identified as performed for the three reference genomes and then classified by motif type (canonical, broad, and degenerated), genomic region (centromere and arm), and continental population group (African (AFR), European (EUR), East Asian (EAS), South Asian (SAS), and American (AMR)).

### Motif conservation visualization

Sequence logos representing nucleotide frequencies across detected CENP-B box motifs were generated in R (v4.4.3) using the ggseqlogo (v0.2)(Wagih 2017) package.

### Scrambled CENP-B box permutation tests

To quantify the specificity and robustness of CENP-B box detection, we performed sequence scramble tests. For example, the broad CENP-B box motif YTTCGNNNNANRCGGGN contains two degenerate positions (Y = C/T, R = A/G) and 8 variable positions (N). We first enumerated the four concrete 17-mers obtained by expanding Y and R (2 × 2 = 4 combinations). For each concrete motif, we consider only the fixed (non-N) positions and treat their bases as a multiset of size k = 11. The number of distinct scrambles for that concrete motif is:

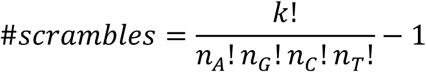

where *n* is the numbers of base in the motif, and the one the same as CENP-B box motif is excluded. The total size of final scrambled motif: 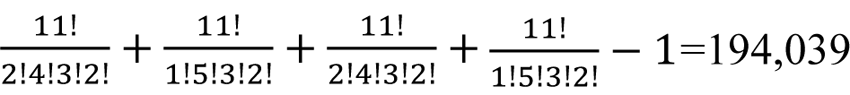. And for the other two motifs, YTTCGTTGGAARCGGGA: 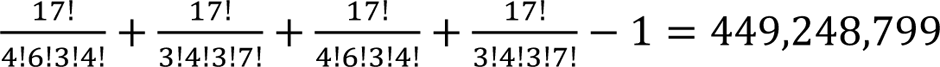; 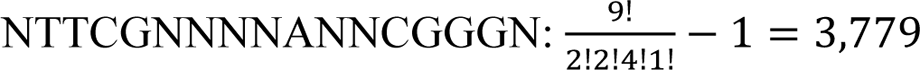 We generated the scramble sequence lists for broad and degenerated motifs and scanned these sequences with three reference genomes.

### Annotation to genomic features

CENP-B box motif coordinates and scrambled-sequence match coordinates located on chromosome arms were annotated to genomic features in CHM13v2.0 using ChIPseeker (v1.42.0) (Yu et al. 2015), in which we classified motif occurrences relative to promoters (±2kb of TSS), exons, introns, downstream regions, and distal intergenic regions.

### ENCODE candidate cis-regulatory elements (cCRE) enrichment analysis

To assess whether the occurrence of CENP-B box motifs on chromosome arms within specific genomic features deviates from randomness, enrichment of CENP-B box motifs within ENCODE candidate cis-regulatory elements (cCREs) (Moore et al. 2026) was further assessed. ENCODE cCREs were classified based on both biochemical signals (chromatin accessibility, H3K4me3, H3K27ac, CTCF and transcription factor) and distance from annotated TSSs (Moore et al. 2026). Enrichment of ectopic motifs within ENCODE cCREs was assessed by permutation testing using 1,000 genome-wide shuffles (*bedtools shuffle*), preserving chromosome assignment and preventing interval overlap. Fold enrichment was calculated as the observed overlap count divided by the mean overlap count across permutations. Empirical P values were computed as

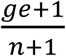

where *ge* is the number of permutations with counts greater than or equal to the observed count and n is the total number of permutations (n = 1000).

### Sequence similarity between CENP-B box and transcription factor binding sites (TFBS)

To evaluate similarity between CENP-B box motif and canonical TFBS motifs, motif-to-motif similarity comparisons were performed using Tomtom (v5.5.9) (https://meme-suite.org/meme/tools/tomtom) against HOCOMOCO human (v11 CORE) database. Tomtom evaluates motif similarity by aligning the position weight matrix (PWM) of a query motif to database motifs across all possible offsets and scoring similarity between aligned columns using the Pearson correlation coefficient (Gupta et al. 2007).

### CUT&RUN CENP-B data processing, alignment and peak calling

- Data preprocessing and alignment

Three CENP-B CUT&RUN biological replicates (SRR9201837, SRR9201839, and SRR9201844) and their corresponding IgG controls (SRR9201847, SRR9201838, and SRR9201846) from (Dumont et al. 2020) were analyzed. Raw sequencing reads were processed using the raw-qc Nextflow pipeline (v3.0.0), which performs sequencing quality control and adapter trimming. Trimmed FASTQ files were then aligned independently to the CHM13 and RPE-1 using BWA-MEM (v0.7.18) (Li and Durbin 2009). Sorted BAM files generated from these alignments were retained for downstream analyses.

- Peak calling

Peak calling was performed by Model-based Analysis of ChIP-Seq (MACS3)(Zhang et al. 2008). Peaks were identified using the following parameters: paired-end mode -f BAMPE, effective genome size -g 3.1e9, --nomodel, --extsize 200, --keep-dup all, -B, and a q-value threshold of -q 0.05. The overlapped peaks among three biological replicates were performed with mergePeaks by homer (v5.1) (Heinz et al. 2010).

- Peak enrichment in CENP-B box

To evaluate peak distribution, bigwigCompare from deepTools (v3.5.6)(Ramírez et al. 2014) was used to compute the log2ratio between the treated sample and the corresponding IgG control. Signal enrichment around CENP-B box motifs was then quantified using computeMatrix from deepTools within ±1 kb flanking regions.

### RNA sequencing and gene expression data analysis

Human foreskin fibroblast BJ-hTERT cells were transduced with lentivirus expressing Doxycycline-inducible full-length CENP-B tagged with mCherry and sorted by FACS. Doxycycline was used at 100 ng/ml for 3 days in Dulbecco modified minimum essential media (DMEM) GlutaMAX medium (Gibco 61965-059), 20% Medium 199 (Sigma Aldrich M4530) supplemented with 10% FBS. RNA was then extracted with the Quick-RNA Miniprep Kit (cat. # R1054, Zymo Research). RNA sequencing libraries were prepared from 1 µg total RNA using the Illumina Stranded mRNA Prep (ligation-based) protocol and sequenced on an Illumina NovaSeq X (NovaSeqX 1.5B-200, 1U configuration) to generate paired-end 100 bp reads (PE100). Data were processed using an in-house RNA-seq analysis pipeline (v4.2.2)(Servant et al. 2024) implemented in Nextflow (v21.10.6) (Di Tommaso et al. 2017). Reads were aligned to the human reference genome (hg38) using STAR (v2.7.6a) (Dobin et al. 2013) with automatic strandedness detection, and gene-level counts were generated using STAR based on GENCODE v34 annotations (GTF), with protein-coding regions provided as BED12. Quality control and metrics were assessed using Picard (v2.25.3)(Broad Institute 2019), RSeQC (v4.0.0)(Wang et al. 2012), Qualimap (v2.2.2-dev)(Okonechnikov et al. 2016), Preseq (v3.1.1)(Daley and Smith 2013), and deepTools bamCoverage (v3.5.1)(Ramírez et al. 2014), with alignment processing performed using samtools (v1.12)(Li et al. 2009) and bcftools (v1.12)(Danecek et al. 2021), and additional filtering using SnpSift (v4.3t)(Cingolani et al. 2012). Genes with low expression were filtered by retaining those with ≥5 counts in more than 25% of samples. Differential expression analysis was performed using the DESeq2 R package (v 1.44.0)(Love et al. 2014). Gene identifiers were mapped from Ensembl IDs to gene symbols using org.Hs.eg.db (v 3.19.1)(Carlson 2024). Preranked gene set enrichment analysis (GSEA) was performed using clusterProfiler (v 4.12.6) (Yu et al. 2012)with the fgsea algorithm. Genes were ranked by log2 fold change, and duplicate gene symbols were collapsed by retaining the entry with the largest absolute effect size. A custom gene set comprising genes with CENP-B box motifs in promoter regions was tested for enrichment.

## Data and code availability

Three CENP-B CUT&RUN samples (SRR9201837, SRR9201839, and SRR9201844) and their corresponding IgG controls (SRR9201847, SRR9201838, and SRR9201846) could be downloaded from SRA. All custom code and critical bed files could be found from: https://github.com/FachinettiLab/CENPBisforCENTROMERE.

## Acknowledgments

We are grateful to Nicolas Altemose, Catalina Salinas, Annapaola Angrisani and all the members of the Fachinetti lab for critical discussions of this work. We would like to acknowledge the Human Pangenome Reference Consortium (BioProject ID: PRJNA730823) and its funder, the National Human Genome Research Institute (NHGRI).

---

**Figure S1.**
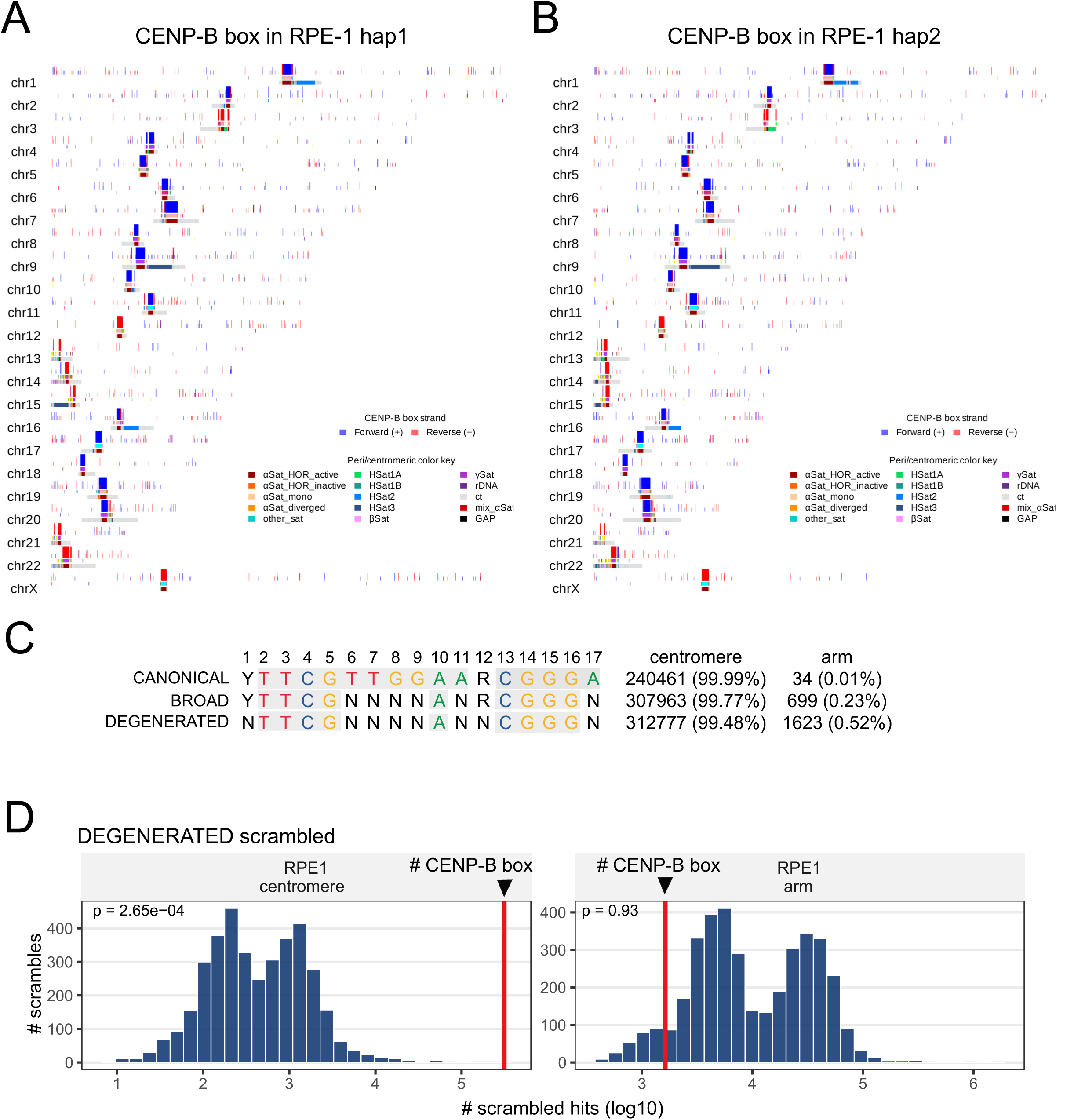
Genome-wide distribution of CENP-B box motifs in the RPE-1_v1.1 reference genome. **(A-B)** Karyoplot of CENP-B box motifs in haplotype 1 **(A)** and haplotype 2 **(B)** in RPE-1_v1.1 reference genome. The first three tracks show CENP-B box motifs based on canonical, broad, and degenerated definitions, with forward- and reverse-strand occurrences shown in blue and red, respectively. (Peri-)centromeric satellite classes include active and inactive αSat higher-order repeats (HORs), diverged αSat, and other satellite families such as HSat1–HSat3, β-satellite (βSat), γ-satellite (γSat), rDNA, and centromeric transition (ct) regions; see color key. **(C)** Genome-wide distribution and total counts of canonical, broad and degenerate CENP-B box motifs in the RPE-1 reference genome. **(D)** Comparison between observed CENP-B box motif counts and scrambled-control motif counts in RPE-1 centromeric regions and chromosome arms. Histograms show the null distributions generated from scrambled motifs, with the observed CENP-B box count indicated by a red vertical line. p-values represent empirical one-sided permutation probabilities, defined as the proportion of scrambled sequences yielding motif counts greater than the observed counts.

**Figure S2.**
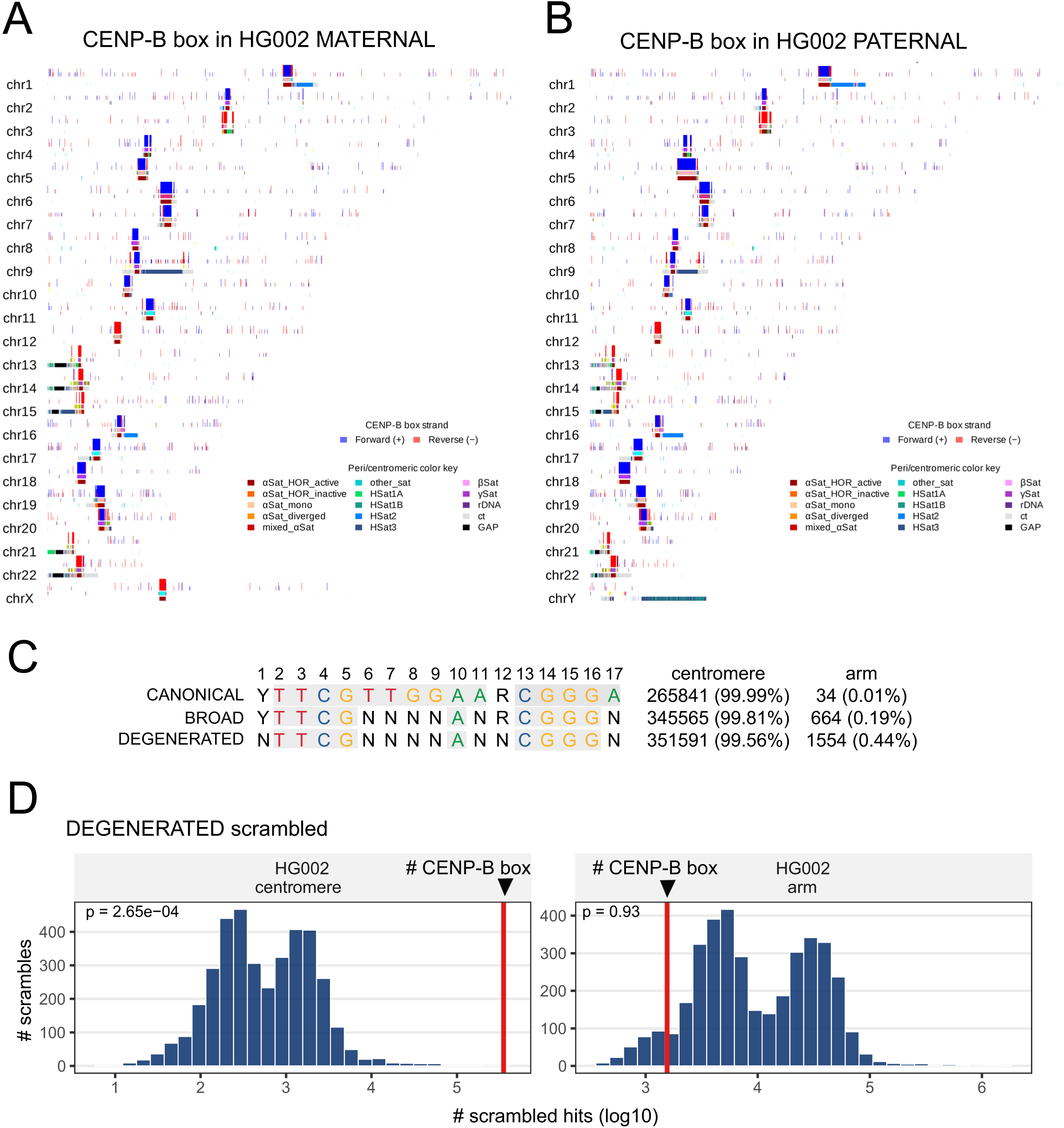
Genome-wide distribution of CENP-B box motifs in the HG002 reference genome. **(A-B)** Karyoplot of CENP-B box motifs in haplotype 1 **(A)** and haplotype 2 **(B)** in HG002 reference genome. The first three tracks show CENP-B box motifs based on canonical, broad, and degenerated definitions, with forward- and reverse-strand occurrences shown in blue and red, respectively. (Peri-)centromeric satellite classes include active and inactive αSat higher-order repeats (HORs), diverged αSat, and other satellite families such as HSat1–HSat3, β-satellite (βSat), γ-satellite (γSat), rDNA, and centromeric transition (ct) regions; see color key. **(C)** Genome-wide distribution and total counts of canonical, broad and degenerate CENP-B box motifs in the HG002 reference genome. **(D)** Comparison between observed CENP-B box motif counts and scrambled-control motif counts in HG002 centromeric regions and chromosome arms. Histograms show the null distributions generated from scrambled motifs, with the observed CENP-B box count indicated by a red vertical line. p-values represent empirical one-sided permutation probabilities, defined as the proportion of scrambled sequences yielding motif counts greater than the observed counts.

**Figure S3.**
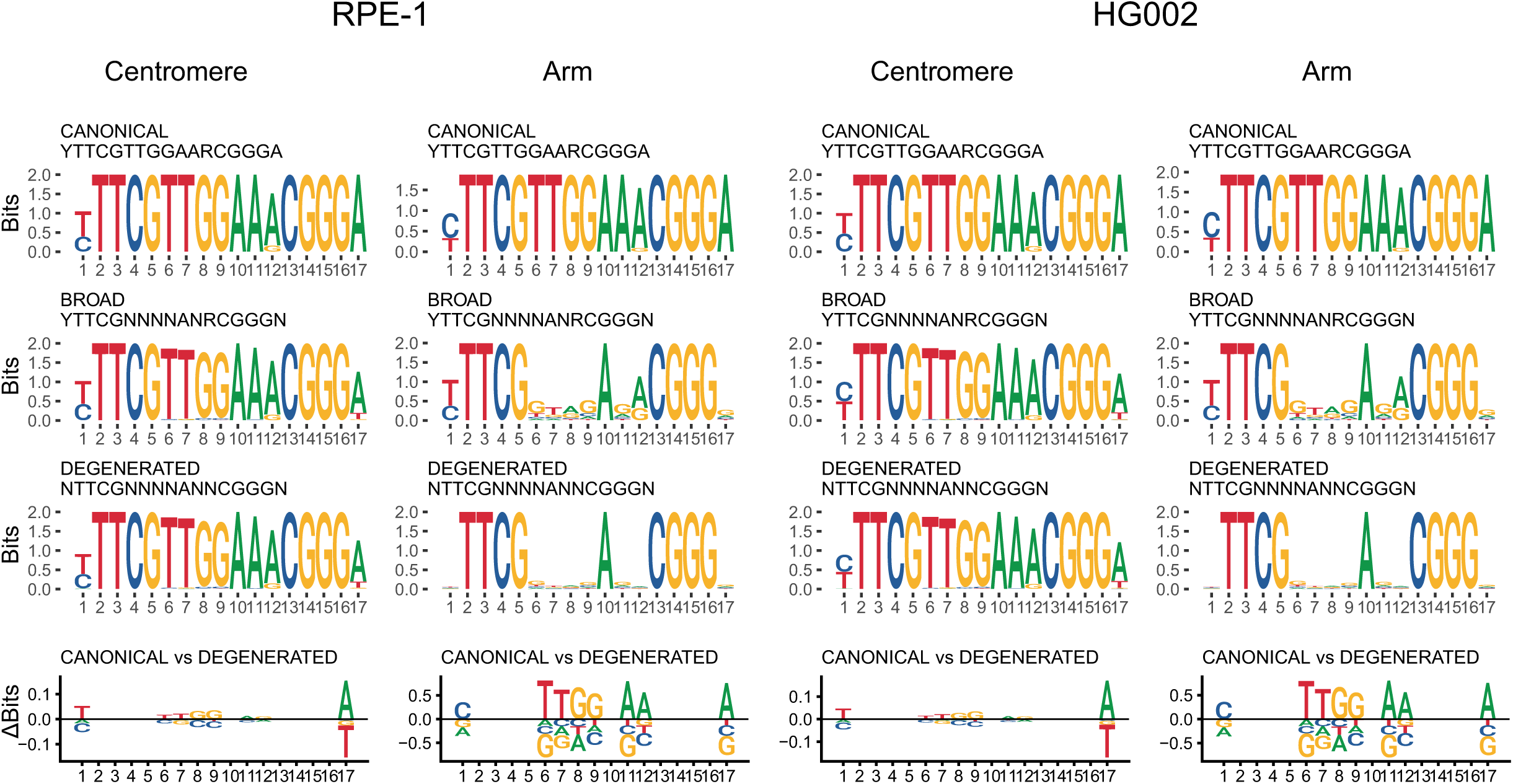
Sequence conservation of CENP-B box motifs in RPE-1 and HG002 reference genomes. Sequence logos illustrating the consensus composition of CENP-B box motifs across human genome assemblies. Three motif definitions are shown: canonical, broad and degenerated. ΔBits plots represent positional differences in nucleotide frequencies between the canonical and degenerated motifs.

**Figure S4.**
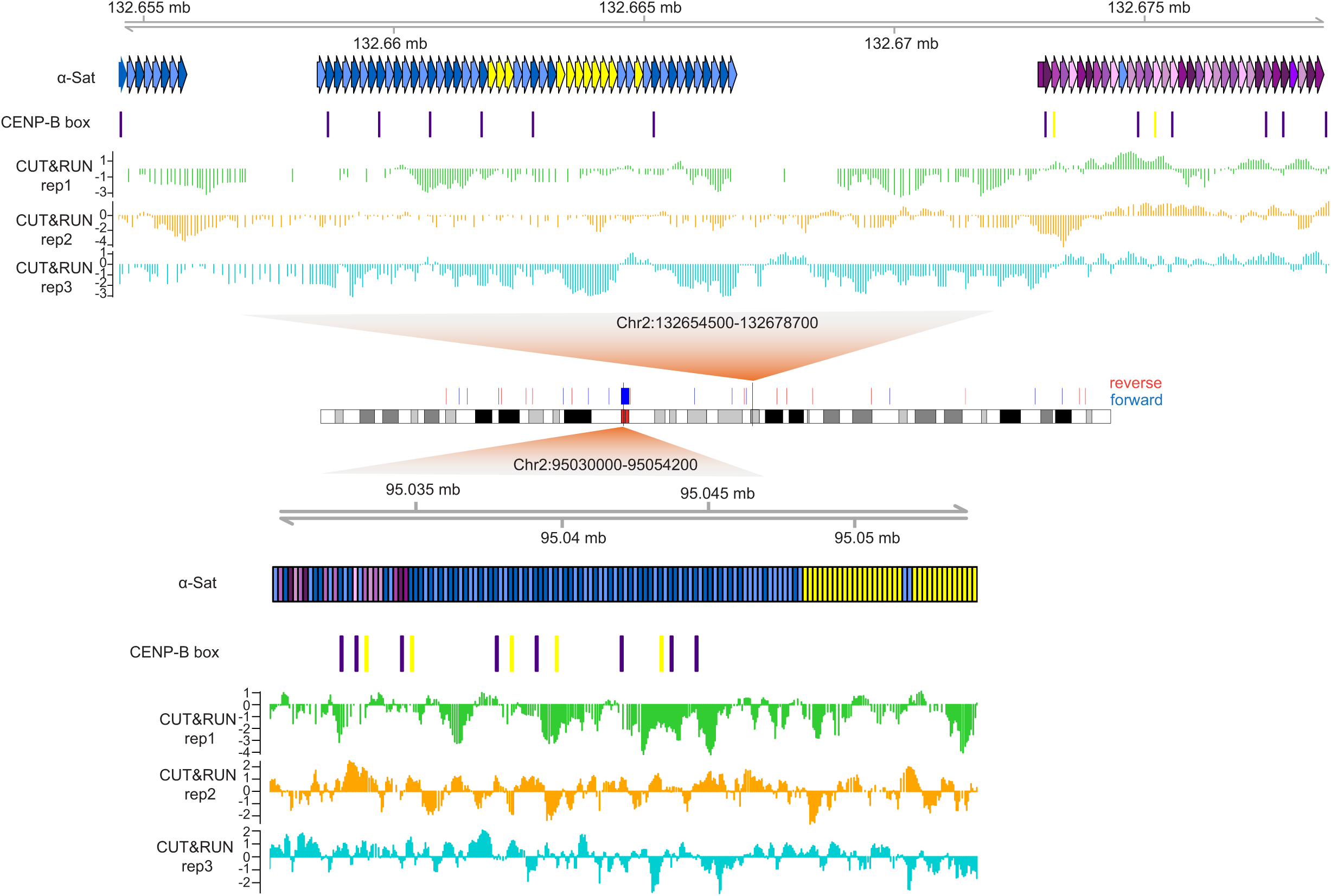
Detection of CENP-B peaks on CENP-B box motifs in centromeric and arm regions of chromosome 2 in CHM13 reference genome. Representative examples of CENP-B box clusters located on the chromosome arms of chromosomes 2 in CHM13 reference genome (top) or at centromeres (bottom). Arm loci coincide with αSat blocks. The genomic locations, orientation, and density of individual CENP-B box motifs, together with CUT&RUN signal tracks across three biological replicates are shown.

**Figure S5.**
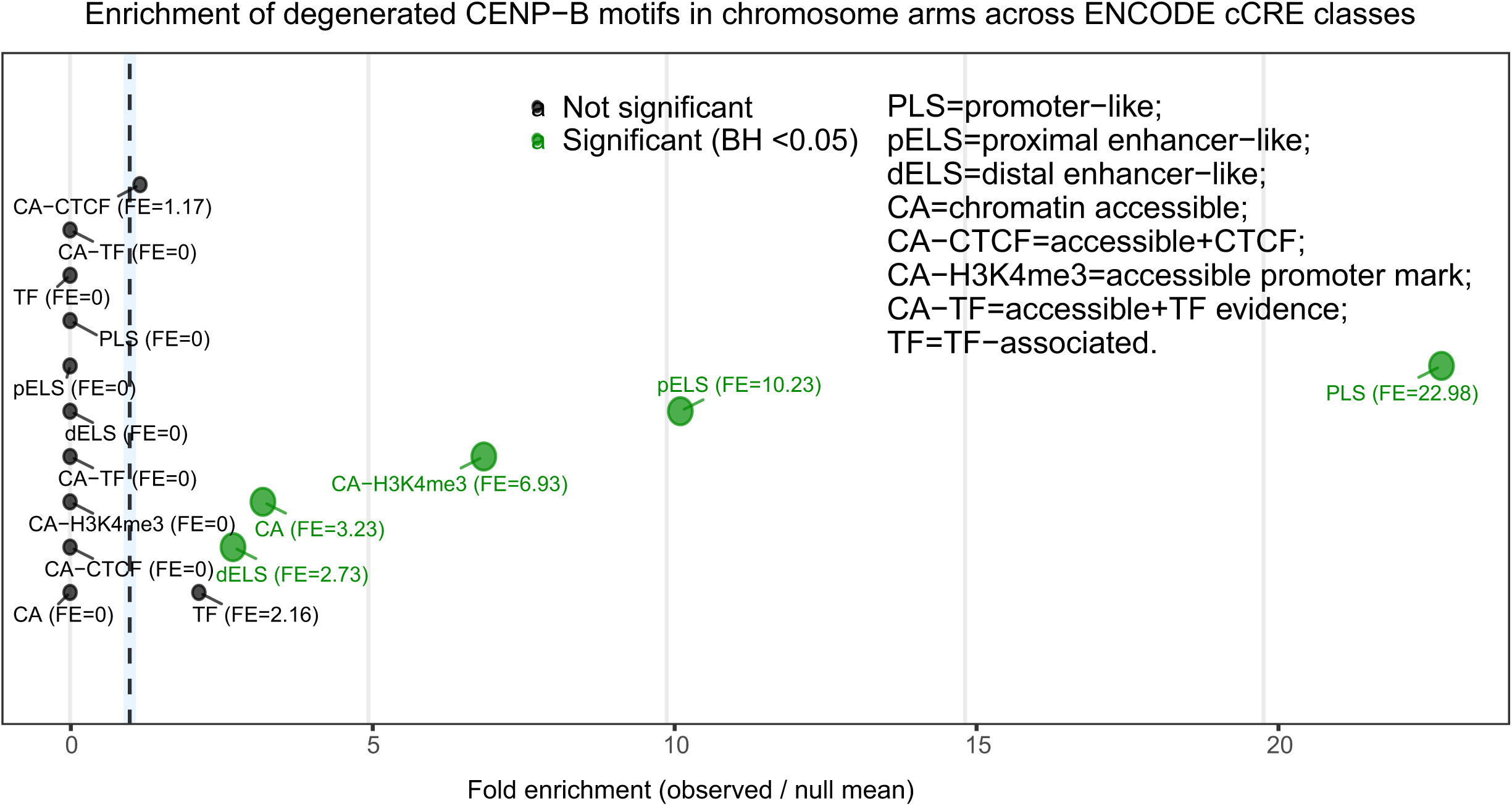
ENCODE cCRE enrichment analysis of CENP-B box motifs on chromosome arms. Enrichment analysis of CENP-B box motifs on chromosome arms relative to ENCODE candidate cis-regulatory element (cCRE) classes. Points indicate observed-to-expected fold enrichment (FE) for each class, with the dashed vertical line marking no enrichment (FE = 1). cCRE classes are defined as follows: promoter-like signatures (PLS), regions with strong promoter-associated chromatin marks; proximal enhancer-like signatures (pELS), enhancer elements located near transcription start sites; distal enhancer-like signatures (dELS), enhancer elements located distal to promoters; CA–H3K4me3, chromatin-accessible regions marked by H3K4me3 and associated with active promoters; CA–CTCF, chromatin-accessible regions bound by CTCF; CA–TF, chromatin-accessible regions with transcription factor binding evidence; and CA, general chromatin-accessible regions without additional specific annotation.

**Figure S6.**
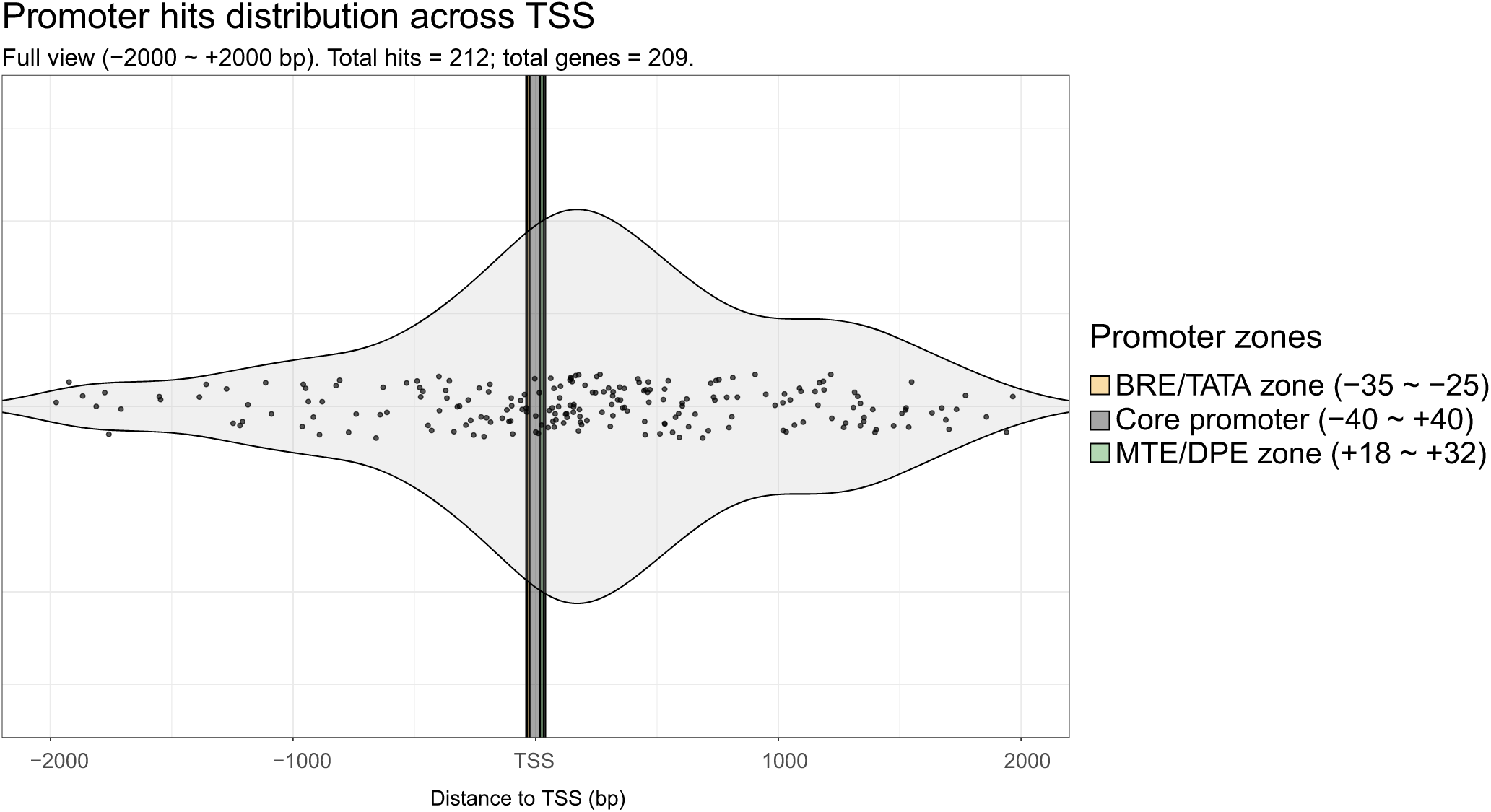
Distribution of CENP-B box motifs relative to transcription start sites (TSS). Violin plot showing the distribution of distances of degenerated CENP-B box motifs to transcription start sites (TSS) within promoter regions in the CHM13 reference genome. Individual motif occurrences are overlaid as jittered points. The analysis includes all motif hits annotated as promoters within a ±2 kb window centered on the TSS (0 bp). The core promoter (−40 to +40 bp) encompasses the minimal region required for transcription initiation. Within this region, the BRE/TATA region (−35 to −25 bp) includes sequence elements involved in transcription factor binding and pre-initiation complex assembly. Downstream of the TSS, the MTE/DPE region (+18 to +32 bp) represents core promoter elements that contribute to transcription initiation in TATA-less promoters.

**Figure S7.**
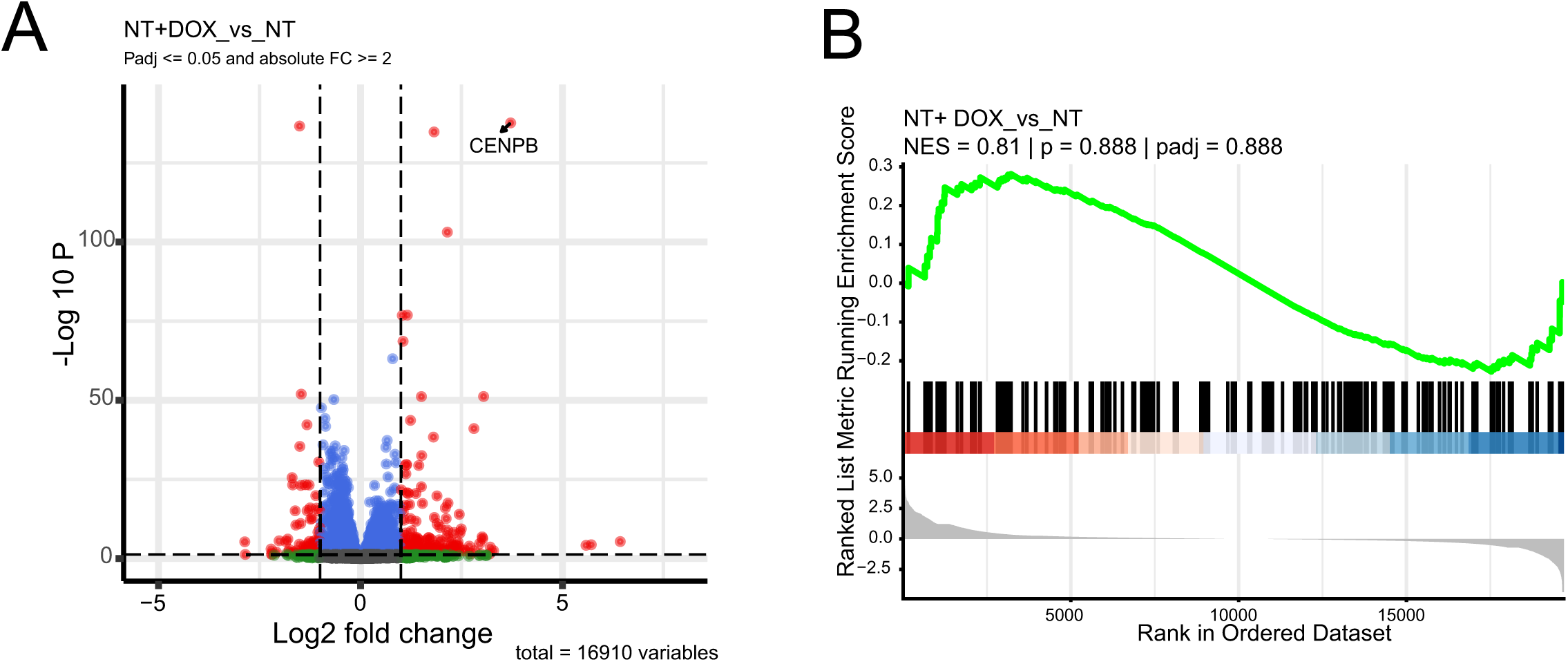
Transcriptomic response to CENP-B overexpression for genes with promoter-associated CENP-B box motifs. **(A)** Volcano plot of differential gene expression in cells overexpressing CENP-B. Genes meeting significance thresholds (adjusted P < 0.05 and absolute value of log2 fold change > 1) are colored in red, and CENPB is highlighted. **(B)** Gene set enrichment analysis of genes containing degenerated CENP-B box motifs within promoter regions (±2 kb from TSS). Genes were ranked by log2 fold change from the differential expression analysis. Normalized enrichment score (NES) and p-value for enrichment significance estimated by fgsea are shown.

**Figure S8.**
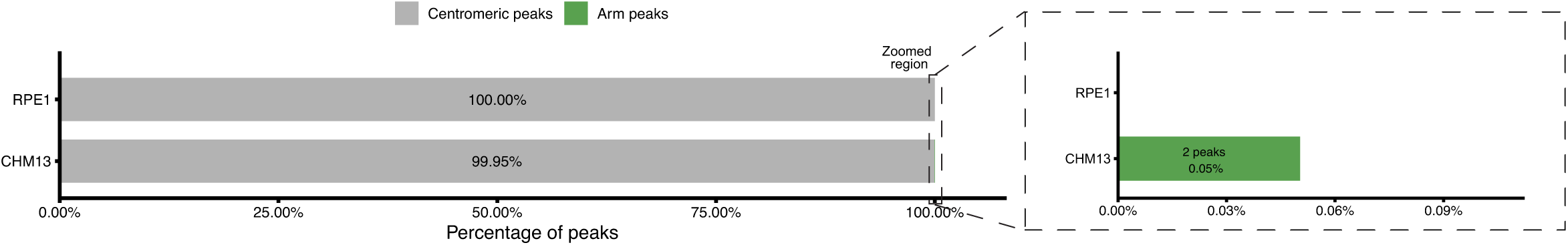
CENP-B CUT&RUN peaks are almost exclusively localized within centromeric CENP-B box motifs. Bar plots show the proportion of CENP-B CUT&RUN peaks overlapping degenerated CENP-B box motifs when mapped on the CHM13 and RPE-1 genome assemblies. Peaks are stratified by genomic location into centromeric regions (grey) and chromosome arms (green).

**Figure S9.**
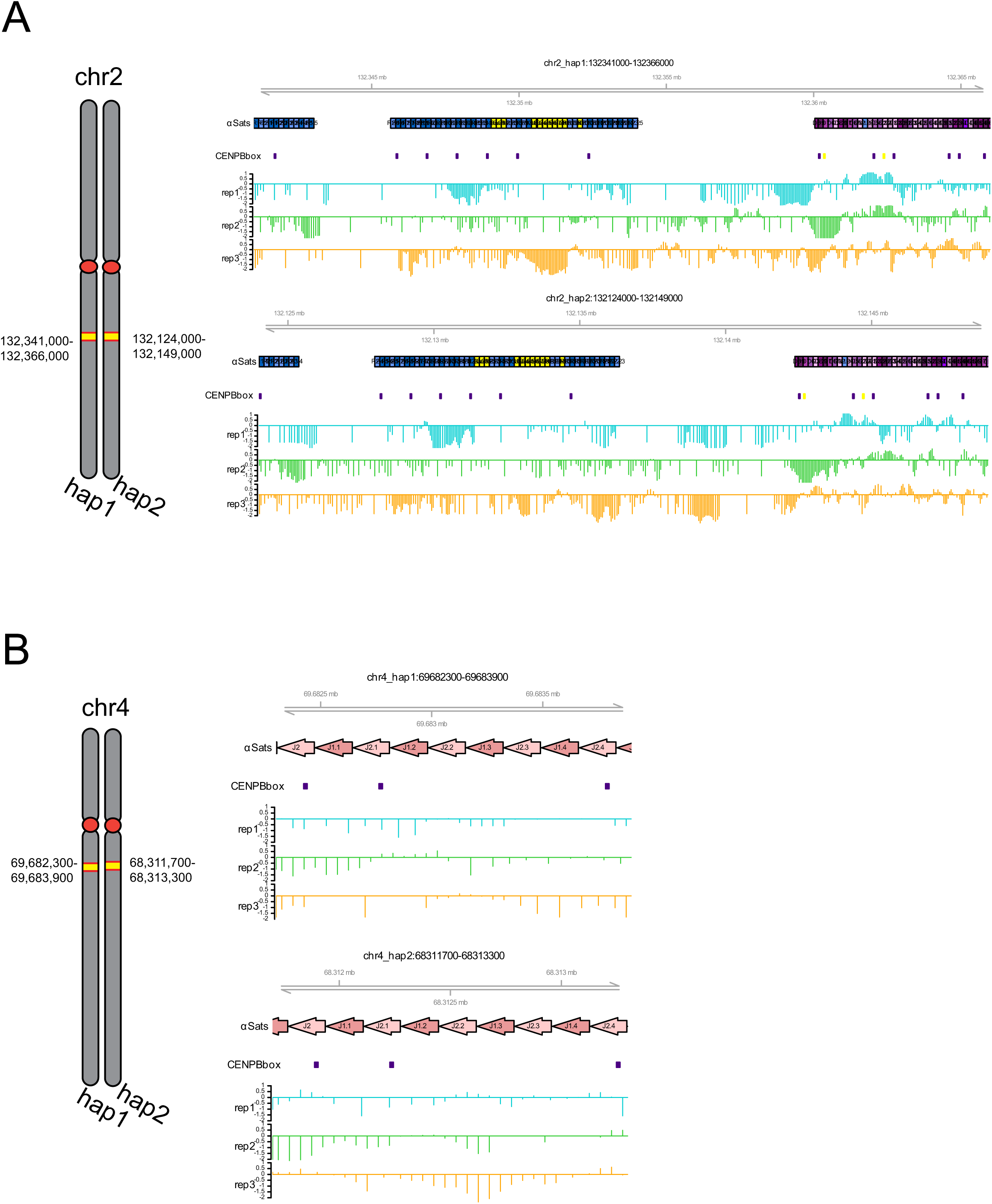
Detection of CENP-B CUT&RUN peaks on CENP-B box motifs in centromeric and arm regions of chromosomes 2 and 4 in the RPE-1 reference genome (**A, B**) Chromosome 2 (A) and chromosome 4 (B) examples. Tracks show the genomic location, orientation, and density of individual CENP-B box motifs, with αSat annotation and CUT&RUN signal across three biological replicates.

**Figure S10.**
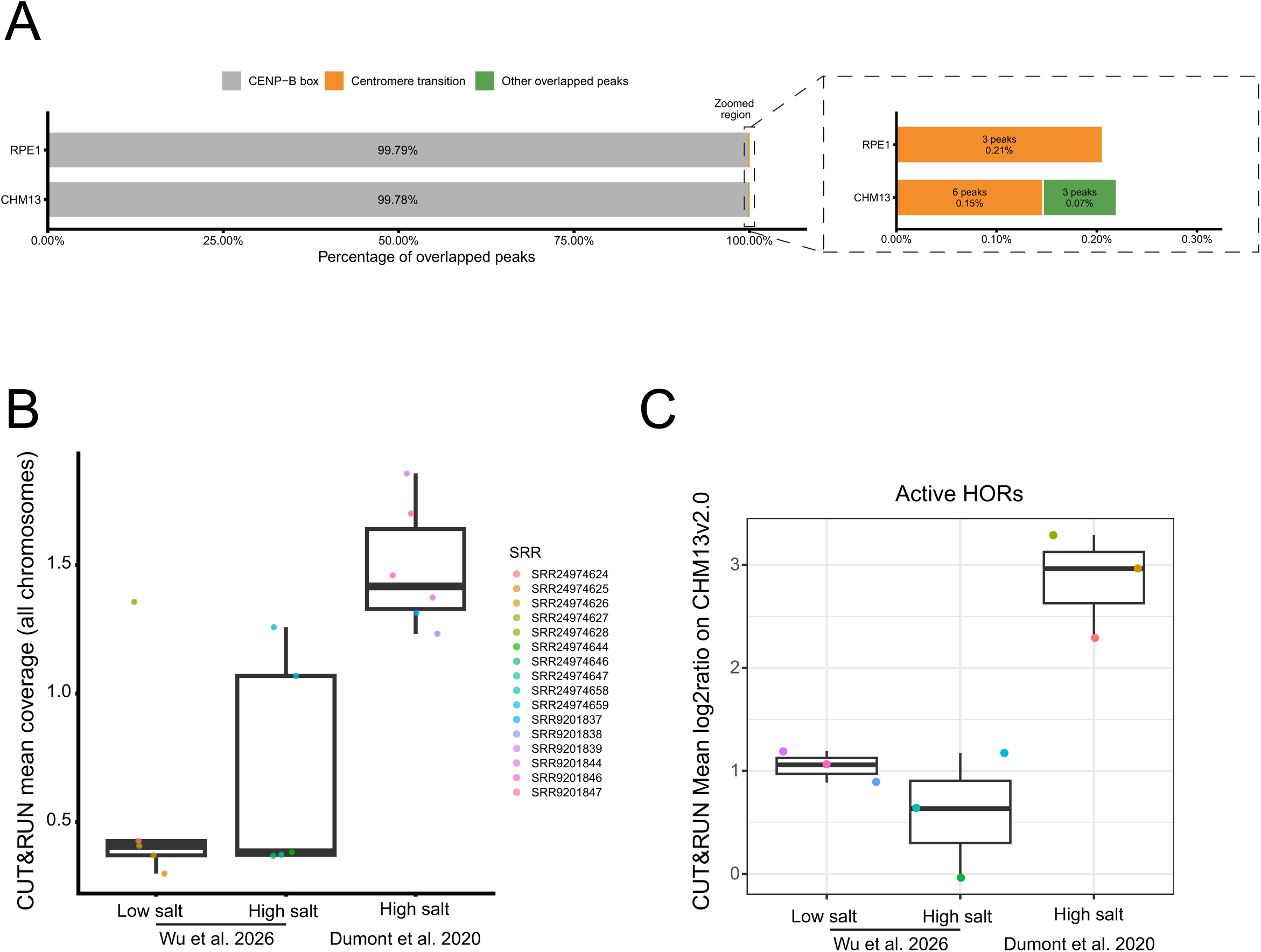
Proportion and localization of CUT&RUN peaks mapped on CHM13 and RPE-1 references, mean coverage, and mean log2 ratio. (A) Bar plots show the proportion of CENP-B CUT&RUN peaks distribution at different genomic locations when mapped on the CHM13 and RPE-1 genome assemblies. Peaks are stratified by genomic location into CENP-B box (gray), centromeric transition (orange), and other overlapped peaks (green). (B) Mean CUT&RUN coverage measured by *mosdepth* for three experimental groups: Wu et al. 2026 low-salt CUT&RUN (rep1: SRR24974629, SRR24974659; rep2: SRR24974627, SRR24974647; and rep3: SRR24974625, SRR24974645); Wu et al. 2026 high-salt CUT&RUN (rep1: SRR24974628, SRR24974658; rep2: SRR24974626, SRR24974646; and rep3: SRR24974624, SRR24974644), and Dumont et al. 2020 high-salt CUT&RUN (rep1: SRR9201837, SRR9201847; rep2: SRR9201839, SRR9201838; rep3: SRR9201844, SRR9201846). (C) Mean log2 CUT&RUN coverage ratio of active HOR regions relative to CHM13v2.0, showing enrichment across the same experimental groups as in B. Jittered points represent individual replicates. All analyses in panels B and C were performed after mapping to CHM13v2.0.

**Figure S11.**
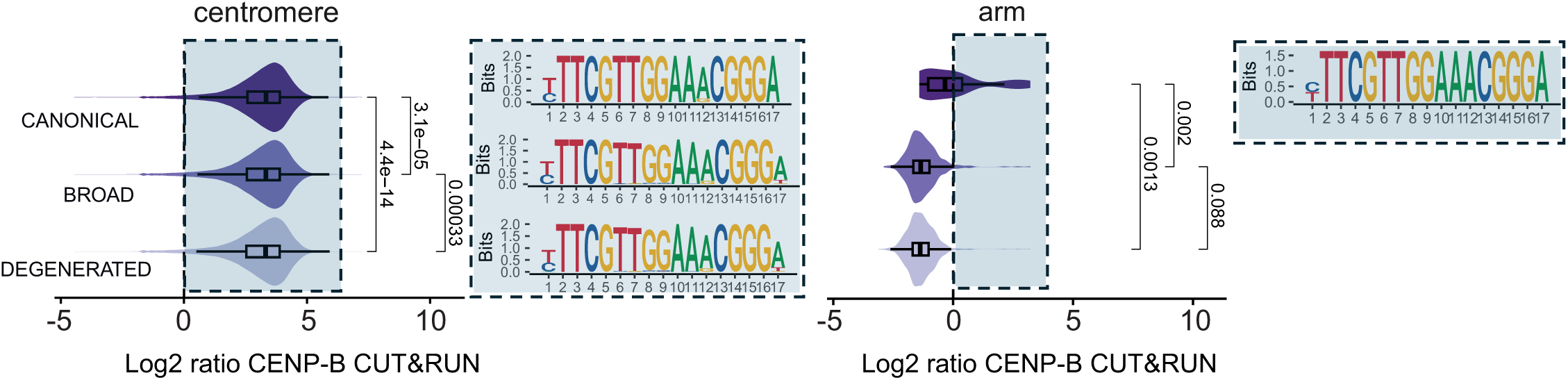
CENP-B CUT&RUN enrichment across CENP-B box motif classes in the CHM13 reference genome. Violin plots show the distribution of log2 CENP-B CUT&RUN enrichment relative to matched IgG control for canonical, broad, and degenerated CENP-B box motifs in centromeric regions and chromosome arms of the CHM13 reference genome. Sequence logos summarize the nucleotide composition of motif instances in each class. P-values indicate comparisons between motif classes using Student’s t-test.

**Table S1.**
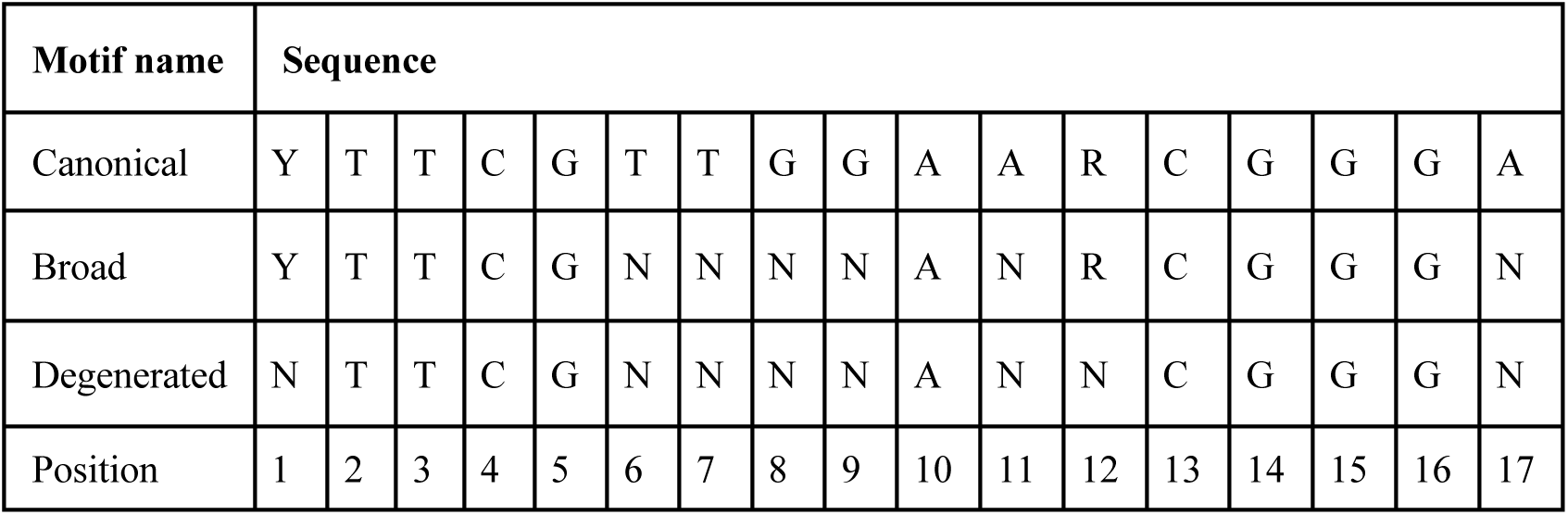
Consensus variants of the human CENP-B box motif used for genome-wide motif searches. Sequences represent the three consensus definitions of the 17-bp CENP-B box motif. Degenerate nucleotide codes follow the IUPAC convention (Y = C/T, R = A/G, N = any nucleotide). Positions correspond to the 17-bp CENP-B box motif.

**Table S2.** Genome-wide distribution of CENP-B box motif variants across human reference assemblies. The table summarizes the total number of CENP-B box motifs detected using three consensus definitions (canonical, broad, and degenerate) across the CHM13, RPE-1, and HG002 genome assemblies. For each motif class, the total number of motif occurrences identified in the genome is reported together with the number and proportion located outside annotated centromeric regions (chromosome arms).

| Reference | Motif name | Total B-box detected | Chromosome arms | Chromosome arms (%) |
| --- | --- | --- | --- | --- |
| CHM13 | Canonical | 127431 | 17 | 0.01% |
|  | Broad | 162532 | 338 | 0.2% |
|  | Degenerated | 166059 | 798 | 0.5% |
| RPE-1 | Canonical | 240495 | 34 | 0.01% |
|  | Broad | 308662 | 699 | 0.2% |
|  | Degenerated | 314400 | 1623 | 0.5% |
| HG002 | Canonical | 265875 | 34 | 0.01% |
|  | Broad | 346229 | 664 | 0.2% |
|  | Degenerated | 353145 | 1554 | 0.44% |

**Table S3.** Summary statistics of scrambled hits across centromeric and chromosome arm regions. The table reports summary statistics for the number of motifs matches per scrambled sequence for both degenerated and broad CENP-B box motif definitions across CHM13 and RPE-1 genome assemblies. Values are provided separately for centromeric and chromosome arm regions and include mean, median, standard deviation, and range of match counts. P detected from empirical permutation test.

| Genome | Term | Broad |  |  | Degenerated |  |  |
| --- | --- | --- | --- | --- | --- | --- | --- |
|  |  | *Centromere* | *Arms* | *Total* | *Centromere* | *Arms* | *Total* |
| CHM13 | Observed | 162190 | 342 | 162532 | 165261 | 798 | 166059 |
|  | Scramble | 170 | 2911 | 3081 | 642 | 10883 | 11526 |
| | Mean<br>(± SD) | (±1307) | (±11937) | (±12443) | (±2174) | (±26848) | (±27970) |
|  | P_greater | 1.03e-5 | 0.86 | 8.04e-4 | 2.65e-4 | 0.94 | 3.44e-3 |
|  | P_twosided | 2.07e-5 | 0.29 | 1.61e-3 | 5.29e-4 | 0.13 | 6.88e-3 |
| RPE-1 | Observed | 307963 | 699 | 308662 | 312777 | 1622 | 314399 |
|  | Scramble | 279 | 5219 | 5498 | 1138 | 21607 | 22744 |
| | Mean<br>(± SD) | (±2193) | (±20420) | (±21222) | (±3803) | (±53593) | (±55600) |
|  | P_greater | 5.17e-6 | 0.84 | 7.37e-4 | 2.65e-4 | 0.93 | 3.70e-3 |
|  | P_twosided | 1.03e-5 | 0.31 | 1.47e-3 | 5.29e-4 | 0.14 | 7.41e-3 |
| HG002 | Observed | 345565 | 664 | 346229 | 351591 | 1554 | 353145 |
|  | Scramble | 344 | 5077 | 5421 | 1413 | 20884 | 22297 |
| | Mean<br>(± SD) | (± 2591) | (± 20035) | (± 21057) | (± 4533) | (± 51795) | (± 54303) |
|  | P_greater | 5.16e-6 | 0.84 | 5.72e-4 | 2.65e-4 | 0.93 | 2.91e-3 |
|  | P_twosided | 1.03e-5 | 0.32 | 1.14e-3 | 5.29e-4 | 0.14 | 5.82e-3 |

**Table S4.**
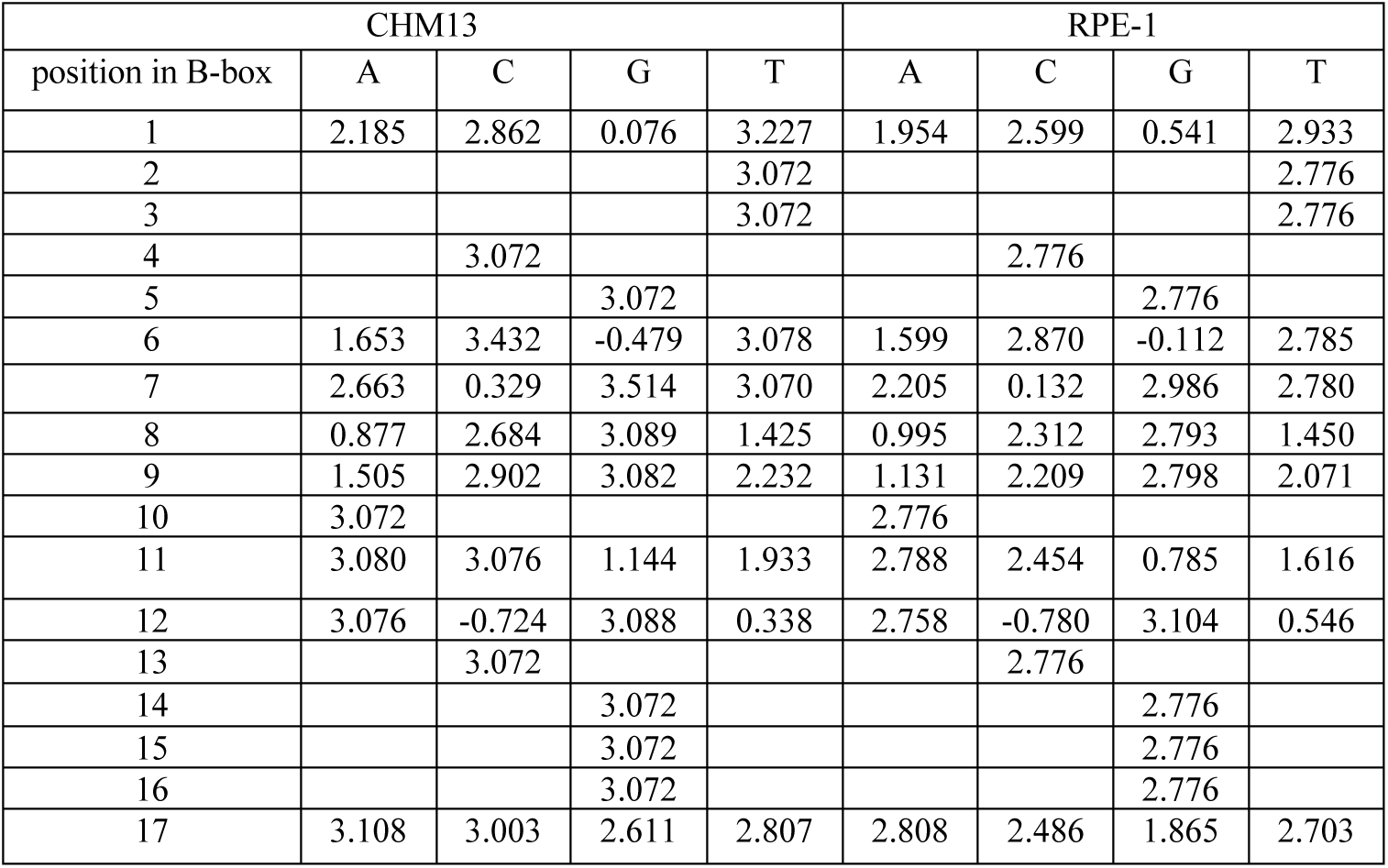
CENP-B CUT&RUN enrichment across the CENP-B box motif. Position-specific nucleotide enrichment across the 17-bp CENP-B box motif calculated from CENP-B CUT&RUN data mapped to the CHM13 and RPE-1 reference genomes. Values represent the log₂ enrichment ratios (CUT&RUN signal relative to IgG control) for each nucleotide (A, C, G, T) at every position of the CENP-B box motif. Higher values indicate stronger preferential enrichment of the CUT&RUN signal associated with that nucleotide at the corresponding motif position, reflecting sequence features that correlate with CENP-B binding.

**Table S5.**
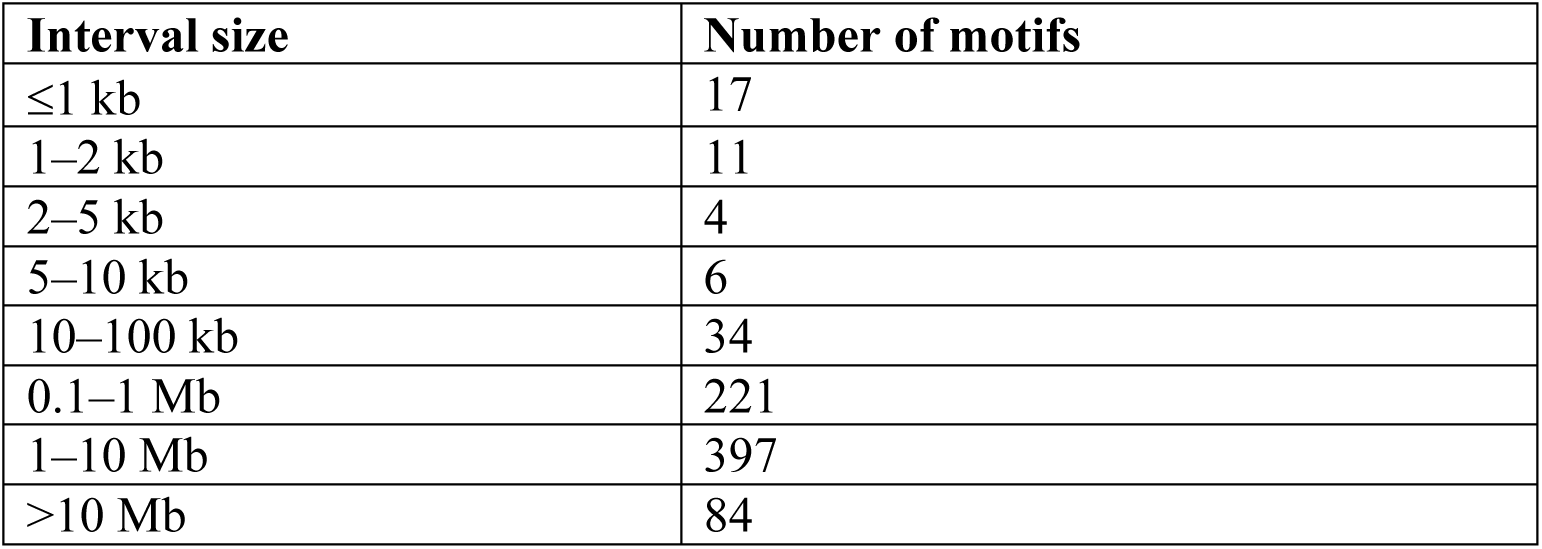
Distance between degenerated motifs within chromosome arms in the CHM13 reference genome.

| Interval size | Number of motifs |
| --- | --- |
| ≤1 kb | 17 |
| 1–2 kb | 11 |
| 2–5 kb | 4 |
| 5–10 kb | 6 |
| 10–100 kb | 34 |
| 0.1–1 Mb | 221 |
| 1–10 Mb | 397 |
| >10 Mb | 84 |

